# PTPRS is a novel marker for early tau pathology and synaptic integrity in Alzheimer’s disease

**DOI:** 10.1101/2024.05.12.593733

**Authors:** Alexandre Poirier, Cynthia Picard, Anne Labonté, Isabelle Aubry, Daniel Auld, Henrik Zetterberg, Kaj Blennow, the PREVENT-AD research group, Michel L. Tremblay, Judes Poirier

**Affiliations:** Division of Experimental Medicine, Faculty of Medicine and Health Science, McGill University, Montréal, Québec, Canada; Goodman Cancer Institute, McGill University, Montréal, Canada; Douglas Mental Health University Institute, Montréal, Québec, Canada; Centre for the Studies in the Prevention of Alzheimer’s Disease, Montréal, Québec, Canada; McGill University, Montréal, Québec, Canada; Victor Phillip Dahdaleh Institute of Genomic Medicine, McGill University, Montréal, Québec, Canada; Department of Psychiatry and Neurochemistry, Institute of Neuroscience and Physiology, The Sahlgrenska Academy, University of Gothenburg, Gothenburg, Sweden; Clinical Neurochemistry Laboratory, Sahlgrenska University Hospital, Mölndal, Sweden; Department of Neurodegenerative Disease, UCL Institute of Neurology, London, United Kingdom; UK Dementia Research Institute at UCL, London, United Kingdom; Hong Kong Center for Neurodegenerative Diseases, Clear Water Bay, Hong Kong, P.R. China; Wisconsin Alzheimer’s Disease Research Center, University of Wisconsin School of Medicine and Public Health, University of Wisconsin-Madison, Madison, WI, USA; University of Science and Technology of China, Hefei, Anhui, P.R. China; Institut du Cerveau et de la Moelle épinière (ICM), Pitié-Salpêtrière Hospital, Sorbonne Université, Paris, France; Department of Biochemistry, McGill University, Montréal, Canada

**Keywords:** Alzheimer’s disease, protein-tyrosine phosphatase receptors, cerebrospinal fluid, tau pathology, synaptic markers, autophagy

## Abstract

We examined the role of protein tyrosine phosphatase receptor sigma (PTPRS) in the context of Alzheimer’s disease and synaptic integrity. Publicly available datasets (BRAINEAC, ROSMAP, ADC1) and a cohort of asymptomatic but “at risk” individuals (PREVENT-AD) were used to explore the relationship between PTPRS and various Alzheimer’s disease biomarkers. We identified that PTPRS rs10415488 variant C shows features of neuroprotection against early tau pathology and synaptic degeneration in Alzheimer’s disease. This single nucleotide polymorphism correlated with higher PTPRS transcript abundance and lower P-tau181 and GAP-43 levels in the CSF. In the brain, PTPRS protein abundance was significantly correlated with the quantity of two markers of synaptic integrity: SNAP25 and SYT-1. We also found the presence of sexual dimorphism for PTPRS, with higher CSF concentrations in males than females. Male carriers for variant C were found to have a 10-month delay in the onset of AD. We thus conclude that PTPRS acts as a neuroprotective receptor in Alzheimer’s disease. Its protective effect is most important in males, in whom it postpones the age of onset of the disease.

## Introduction

Synaptic dysfunction is a central pathologic feature of Alzheimer’s disease (AD), with synaptic loss preceding neuronal loss in specific brain regions (1). Synaptic plasticity and neuroplasticity result from the complex integration of cellular proliferation, differentiation, migration, and axon guidance of neural cells. At the molecular level, these elaborate processes depend on interactions between cellular receptors associated with internal downstream communication pathways. Such receptors include cadherins, Ig superfamily proteins, neurexins, neuroligins, and leucine-rich repeat proteins (LRRPs), as well as receptor tyrosine kinases (RTKs) and receptor protein-tyrosine phosphatases (PTPRs). PTPRs enzymes are encoded by 47 different genes subdivided into eight sub-groups based on their general conserved structures and homology. The R2B sub-group is particularly interesting for AD research, as they are mainly expressed in the central nervous system (CNS). The R2B subgroup is composed of PTPRD (delta), PTPRS (sigma), and PTPRF (LAR). These enzymes share two intracellular phosphatase segments: an active catalytic phosphatase (D1) and an inactive but regulatory PTPase (D2). Moreover, these three genes also share heterogeneous extracellular segments generated by complex alternative splicing events, leading to Ig-like domains and various fibronectin type-III domains. These three R2B RPTPs are also subject to different proteolytic cleavages that allow the shedding of their extracellular domains. Their ability to regulate cell adhesion and cell signaling creates a balance between synaptic plasticity or arrest (2)

Recently, PTPRD was found to be genetically associated with neurofibrillary tangle accumulation in AD (3). While the detailed molecular mechanism of this association remains unclear, the effect of the PTPRD rs560380 polymorphism on neurofibrillary tangle accumulation is powerful (p = 3.8×10^-8^) and consistent with another report linking the *PTPRD* gene locus to AD dementia risk (4). The delta variant is highly expressed in the brain, where it has been implicated in synaptic differentiation. On the other hand, a null allele in mice leads to memory impairment and altered electrophysiological responses in the hippocampus (5). The second member of the R2B subfamily, PTPRS, is mainly associated with autophagy regulation in neurons (6, 7). Noteworthy, we and others have shown that PTPRS is involved in CNS plasticity, axonal guidance, and other features of neuronal development (6, 8–10). PTPRS is involved in the innervation of excitatory and inhibitory synaptic inputs in cortical areas (11–13). Inhibition of PTPRS is associated with increased neuronal survival, which partly relates to its role as a suppressor of neuronal autophagy (6).

Although no formal genetic association has been reported between the *PTPRS* gene and AD, studies using *Ptprs*-deficient models of Alzheimer’s pathology revealed that neuronal *Ptprs* mediates both amyloid and tau pathogenesis in double transgenic mice by binding to the amyloid precursor protein (*APP*) and interfering with its binding to the beta-secretase, diminishing the APP proteolytic products generated by the beta- and gamma-secretases (14). Tau aggregation, neuroinflammation, synaptic loss, and cognitive deficit all showed clear dependency on the expression of *Ptprs* in the receptor-deficient rodent brain (14).

In the present study, we leveraged the data collected in two large cohorts dealing with AD pathophysiology, namely the pre-symptomatic PREVENT-AD cohort and the ROSMAP cohort, to examine the *PTPR*S locus that displays pleiotropic effects on amyloid, tau, and synaptic pathologies and thus influence multiple insults and pathways leading to neurodegeneration. Such loci could be an excellent target for further investigation in understanding synaptic pathology and exploring the shared pathways associated with brain resilience to different neuropathological processes.

## Materials and Methods

All procedures were approved by the McGill University Faculty of Medicine Institutional Review Board and complied with the ethical principles of the Declaration of Helsinki.

### PREVENT-AD cohort

#### Study participants

PREVENT-AD is an observational cohort of healthy older adults at increased risk of AD dementia (15). PREVENT-AD enrolled more than 400 cognitively unimpaired participants aged 60 years or older having a parent or at least two siblings diagnosed with AD dementia. Participants were followed up annually with structural and functional magnetic resonance imaging (MRI) medical, and cognitive assessments. Participants also gave blood at each visit, and a subset of 160 volunteered for at least one lumbar puncture (LP). More recently, a partially overlapping sample (n= 129) also volunteered for brain positron emission tomography (PET) scans to assess Aβ and *Tau* deposition *in vivo*.

#### CSF measurements

Lumbar punctures were performed using a Sprotte 24-gauge atraumatic needle following an overnight fast. CSF samples were centrifuged within 4 hours to exclude cells and insoluble material. Blood samples are obtained before LPs to ensure a temporal relationship between peripheral and CNS measures. PTPRS protein levels were measured using a specific enzyme-linked immunosorbent assay (ELISA) (human PTPRS ELISA kit, abx152898, Abbexa Ltd., UK). It is important to note that the exact recognition sequence by the PTPRS detection antibody is unavailable. However, the supplier confirms that it is in the “extracellular proximal membrane domain” of the PTPRS protein. The CSF AD biomarkers P-tau, T-tau, and Aβ_42_ were measured using a validated INNOTEST ELISA kits (P-tau, Cat.# 81581; T-tau, Cat.# 81579, and Aβ_42_, Cat.# 81583) from Fujirebio Europe, Ghent, Belgium, following procedures from the BIOMARKAPD consortium (16). Data were collected between September 2011 and August 2017 and archived in PREVENT-AD data release 6.0 (https://preventad.loris.ca/main.php). Immunoprecipitated SNAP-25 and SYT-1 from CSF were quantified by high-resolution selected ion monitoring (HR-SIM) analyses on a quadrupole–orbitrap mass spectrometer Q Exactive as described in Brinkmalm et al. (17) and Öhrfelt et al. (18). CSF neurogranin and GAP-43 concentrations were assessed using validated ELISAs described before (19, 20).

#### Genotyping and Imputation

Automated DNA extraction from buffy coat samples was performed using the QiaSymphony DNA mini kit (Qiagen, Toronto, Canada). Genotypes were determined with the Illumina Infinium Omni2.5M-8 array (Illumina, San Diego, CA, USA). The PLINK toolset (http://pngu.mgh.harvard.edu/purcell/plink/) was used to: 1) filter gender mismatches, 2) filter missingness at both the sample-level (*<* 5%) and SNP-level (*<* 5%), 3) assess sample heterozygosity and 4) filter SNPs in Hardy-Weinberg disequilibrium (p*>*0.001). Only post-imputed SNPs with an info score *>* 0.7 were considered.

#### The Religious Order Study and the Memory and Aging Project (ROSMAP)

The Religious Orders Study (ROS) was established in 1994, and it includes nuns, priests, and brothers from across the United States [26]. The Rush Memory and Aging Project (MAP) started in 1997 and includes laypeople from Illinois. Participants from the cohorts were cognitively normal at enrolment and were followed annually with neuropsychological evaluation and blood tests and consented to genotyping and brain donation [26]. Post-mortem evaluation was performed to assess AD pathology using CERAD and Braak staging. Datasets are available at the NIAGADS repository at https://dss.niagads.org/cohorts/religious-orders-study-memory-and-aging-project-rosmap/.

##### TMT Proteomic Data

340 cortical prefrontal brain tissues from the community-based aging ROSMAP cohort were analyzed by a mass spectrometry-based protein quantification approach using isobaric multiplex tandem mass tags (TMT) as described before by Ping et al. 2018 [27]. TMT labeling with synchronous precursor selection (SPS)-MS3 for reporter ion quantitation was used to achieve comprehensive global quantitation of 100 mg (wet tissue weight) pre-frontal cortex from healthy controls and sAD cases. In total, 127,321 unique peptides were identified from over 1.5 million peptide spectral matches (PSMs), mapped to 11LJ840 unique protein groups; representing 10,230 gene symbols, which map to ∼65% of the protein-coding genes in the brain. Three major isoforms of *PTPRS* expressed in the brain are available in the ROSMAP dataset: the total length (Q13332) wild-type variant and common isoform variants (Q13332-2, Q13332-4 and Q13332-7) which are particularly prevalent in the CNS.

#### Genotyping and Imputation

Imputed genome-wide genotype data from ROSMAP was obtained from the Accelerating Medicines Partnership in Alzheimer’s Disease (AMP-AD) Knowledge Portal (synapse ID: syn3157329). DNA was extracted from blood or post-mortem brain tissue from ROSMAP participants and genotyped on the Illumina OmniQuad Express platform. After quality control (genotype success rate > 0.95, Hardy–Weinberg equilibrium p > 0.001, and mishap test < 1 × 10^9^) and excluding population outliers (participants of non-European Ancestry inferred from the genotype covariance matrix to avoid confounding from population stratification), 382 participants underwent with genome-wide genotyping. Imputation was done on the 1000 Genomes Project (Phase 1b data freeze) reference panel and after removing rare (MAF < 0.01) or poorly imputed variants (INFO score < 0.3). Further details are available through previous publications (De Jager et al., 2018).

#### Alzheimer Disease Center Dataset 1 (ADC1)

The NIA ADC cohorts (1–7) include subjects ascertained and evaluated by the clinical and neuropathology cores of the 39 past and present NIA-funded Alzheimer’s Disease Centers (ADC). Data collection is coordinated by the National Alzheimer’s Coordinating Center (NACC). NACC coordinates collection of phenotype data from the ADCs, cleans all data, coordinates implementation of definitions of AD cases and controls, and coordinates collection of samples. Biological specimens are collected, stored, and distributed by the National Cell Repository for Alzheimer’s Disease (NCRAD). ADC1 datasets are available upon request at https://www.niagads.org/datasets/ng00022. ADC1 was first published in Naj et al. 2012.

#### Human Subjects Demographics and mapping of PTPRS rs10415488

The ADC1 sample set includes 1985 late-onset AD cases and 523 cognitively normal controls which were genotyped using different platforms followed by imputation to generate a common set of 2,324,889 SNPs. Uniform and stringent quality control measures were applied to all datasets to remove low quality and redundant samples and problematic SNPs as per Naj et al., 2012. Dataset NC00022-ADC1 includes information about sex, race, autopsy characteristics, APOE genotype, BRAAK stage, ethnicity and age at onset. We used p-link version 1.9 to extract PTPRS rs10415488 genotype and assessed its impact on age at onset.

### Brain eQTL Almanac (BRAINEAC)

Central nervous system (CNS) tissues originating from 134 cognitively unaffected control individuals were collected by the Medical Research Council Sudden Death Brain and Tissue Bank, Edinburgh, UK,(21) and the Sun Health Research Institute (SHRI), an affiliate of Sun Health Corporation, USA.(22) Anatomical regions of interest were sampled from brain coronal slices on dry ice. A detailed description of the samples used in the study, tissue processing, and dissection is provided in Trabzuni *et al.*(23) All samples had fully informed consent for retrieval and were authorized for ethically approved scientific investigation (Research Ethics Committee number 10/H0716/3).

#### Genotyping

Genomic DNA was extracted from sub-dissected samples of human post-mortem brain tissue using Qiagen’s DNeasy Blood & Tissue Kit (Qiagen, UK). All samples were genotyped on the Illumina Infinium Omni1-Quad BeadChip and on the Immunochip, a custom genotyping array designed for the fine-mapping of auto-immune disorders.(24, 25) After standard quality controls (removal of suspected non-European descent individuals, samples with call rate < 95% and checks on reported sex status, cryptic relatedness, autosomal heterozygosity rate check, monomorphic SNPs or call rate < 95%, no genomic position info or redundant SNPs, p-value for deviation from HWE < 0.0001, genotyping call rate < 95%, less than two heterozygotes present, mismatching alleles 1000G even after allowing for strand), imputation was performed using MaCH.(26, 27) This resulted in ∼5.88 million SNPs and ∼577 thousand indels with good post-imputation quality (Rsq > 0.50) and minor allele frequency of at least 5%.

#### Microarray

Total RNA was isolated from human post-mortem brain tissues based on the single-step method of RNA isolation(28) using the miRNeasy 96 kit (Qiagen). The quality of total RNA was evaluated by the 2100 Bioanalyzer (Agilent) and RNA 6000 Nano Kit (Agilent) before processing with the Ambion® WT Expression Kit and Affymetrix GeneChip Whole Transcript Sense Target Labeling Assay and hybridization to the Affymetrix Exon 1.0 ST Arrays following the manufacturer’s protocols. Further details regarding RNA isolation, quality control, and processing are reported in Trabzuni *et al.*(23) Gene-level expression was estimated for 26 thousand genes by calculating the Winzorised mean (below 10% and above 90%) signal of all probe sets corresponding to each gene. The resulting expression data was adjusted for brain bank, gender, and batch effects in Partek’s Genomics Suite v6.6 (Partek Incorporated, USA).

##### Statistical analyses

We compared PREVENT-AD demographic characteristics of PTPRS rs10415488 C allele-negative (TT) and C-allele-positive (CT/CC), *APOE* ε4-negative and *APOE* ε4 positive unimpaired older adults using Fisher exact or Kruskal-Wallis tests where appropriate. We then tested for associations between CSF AD biomarkers (Aβ_42_, t-*tau*, P-*tau*) with CSF PTPRS using general linear models, adjusted for age and gender and stratified by APOE-ε4 positivity. We also tested for association between CSF PTPRS levels with CSF presynaptic proteins (CAP-43, SYT-1, SNAP-25 using general linear models, controlling for age and sex. Independent t-tests were used to compare ROS-MAP PTPRS mRNA levels as a function of gender, *APOE* ε4 status, CERAD and Braak stages. Finally, we also for the association between cortical PTPRS mRNA levels with presynaptic proteins mRNA prevalence (SYT-1, SNAP-25) using general linear models, controlling for age and sex.

## Results

### Brain *PTPRS cis*-regulation analysis identifies both rs10415488 and *APOE4* as potent modulators of PTPRS mRNA levels in multiple brain regions with no significant effect on tangles or amyloid plaques pathology estimated by Braak and CERAD staging

Cis-eQTL analysis of the *PTPRS* gene region identified a polymorphism that reached locus-wide significance in the BRAINEAC cohort, composed of 134 brains free of neurodegenerative diseases (Fig. 1, A). We further validated this association using the protein abundance of cortical PTPRS in the ROSMAP cohort (Sup. Fig. 1). For the rest of the study, we thus used rs10415488 TT, CT, and CC carriers as a surrogate for low, normal, and higher PTPRS abundance, respectively. We next noted a significant reduction of the cortical levels of PTPRS transcripts in AD versus control subjects in the *APOE4* carriers of the ROSMAP cohort (Fig. 1, B). We did not find, however, any significant associations between PTPRS transcript levels and disease severity, as assessed by CERAD (Fig. 1, C) and Braak staging (Fig. 1, D). These results most likely indicate that the abundance of PTPRS does not impact the later stages of AD. We thus reoriented our study to the pre-symptomatic phase of the disease.

**Fig. 1:**
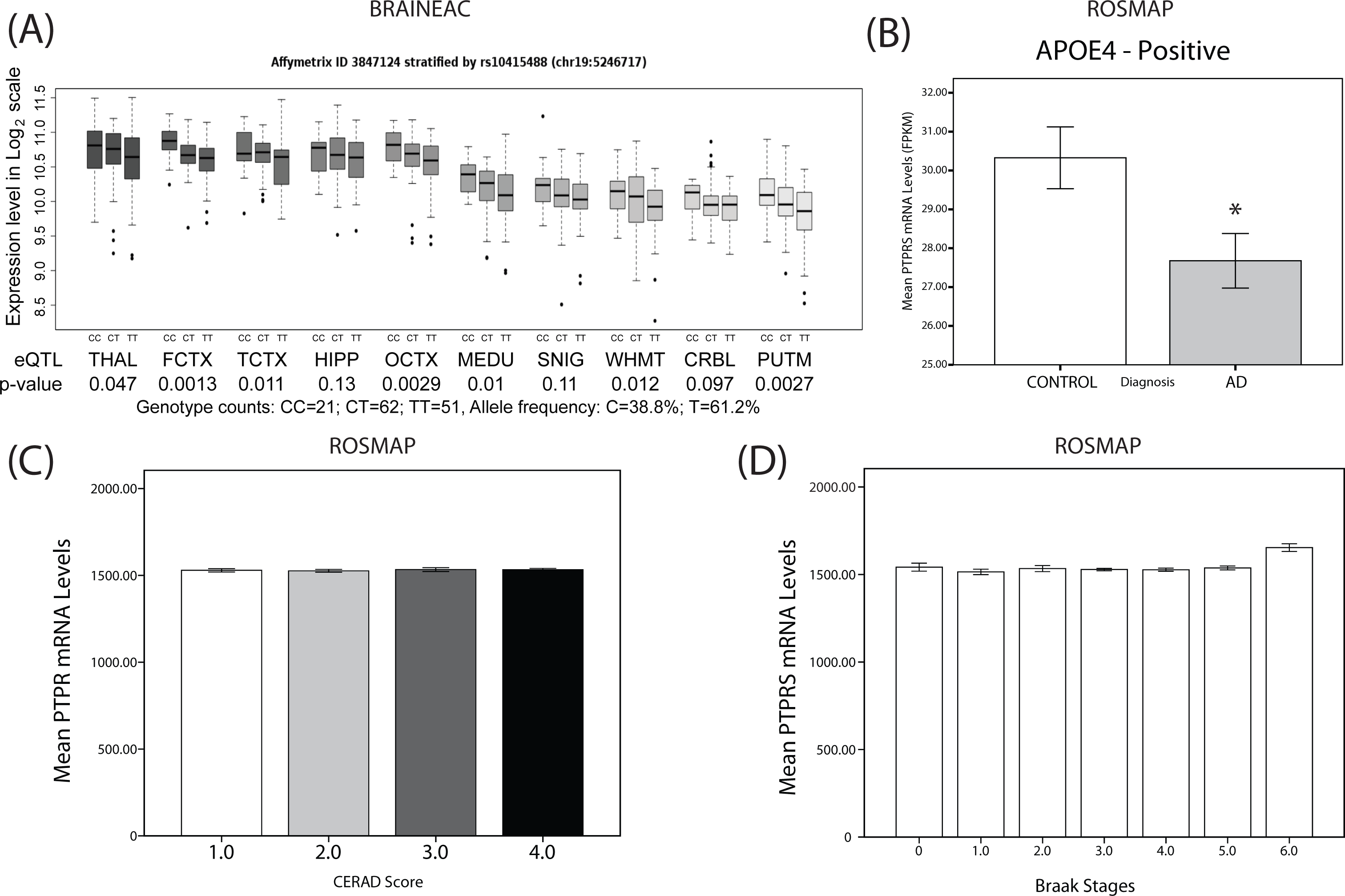
Identification of *PTPRS* rs10415488 variant C as a SNP associated with increased expression. (A) *PTPRS* mRNA levels versus rs10415488 dosage (CC/CT/TT) and brain area from the BRAINEAC database. (B) Mean PTPRS mRNA levels (Fragments Per Kilobase of transcript per Million mapped reads (FPKM)) in controls and AD subjects (*APOE4*-carriers). (C) Mean *PTPRS* mRNA levels in function of CERAD score (1–4). N= 121, 159, 55, 140 for each CERAD score (1–4), respectively. (D) Mean *PTPRS* mRNA levels in the function of Braak stages (0–6). N= 8, 43, 50, 150, 143, 79, and 2 for each Braak stage (0–6) respectively). Statistical analysis: (A) Simple linear regression with Pearson’s correlation. (B) Bilateral unpaired T-test (C-D) One-way ANOVA with Dunnet’s multiple comparison. *p≤0.05, **p≤0.01, ***p≤0.001, ****p≤0.0001.

### Effect of PTPRS rs10415488 C variant on AD disease risk/protection

Next, we investigated the effect of the PTPRS rs10415488 C variant on the overall risk of developing AD using two distinct populations of case/control subjects. A significant association between the C allele, *PTPRS* “high”, and AD risk reduction in the genetically homogeneous Quebec Founding population (QFP) isolate from Eastern Canada (p = 0.02, O.R. 0.86) and in the larger heterogeneous IGAP II GWAS (p = 0.01, O.R. 0.97) was observed (Table I). To validate the implication of PTPRS in AD risk, we performed a mirror analysis, this time on the TT, *PTPRS* “low” variant (Sup. Table I). We found that in the QFP (p=0.008, OR=1.30), TT carriers had an increased risk of developing AD. Upon stratification for *APOE4*, we found that the OR climbed to 3.12 and slightly declined in *APOE4*-positive and -negative individuals, respectively (Sup. Table I). These observations indicate, retrospectively, that sigma phosphatase receptor abundance is linked to the risk of developing AD.

**Table I:**
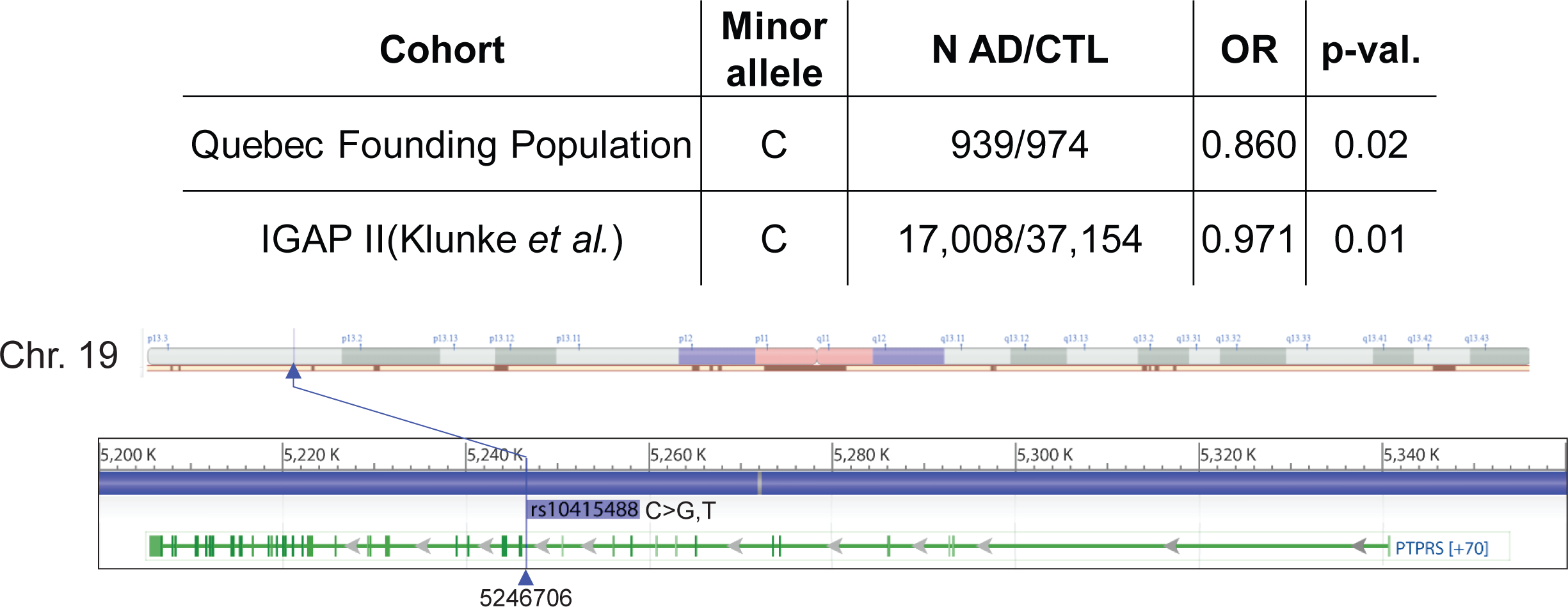
rs10415488 C variant is associated with protection from Alzheimer’s disease. (top) Number of Alzheimer’s patients/control subjects, odds ratio (OR), and p-value of the OR in the Quebec Founding Population (QFP) and IGAP II cohorts. (bottom) Chromosomal localization of rs10415488 in the *PTPRS* locus.

| Cohort | Minor allele | N AD/CTL | OR | p-val. |
| --- | --- | --- | --- | --- |
| Quebec Founding Population | C | 939/974 | 0.860 | 0.02 |
| IGAP II(Klunke *et al.*) | C | 17,008/37,154 | 0.971 | 0.01 |

Chr. 19

### Cortical PTPRS protein isoform levels as a function of SYT1 and SNAP25 protein levels

Using ROSMAP’s TMT protein dataset, we examined the relationship between different cortical PTPRS protein isoforms and two synaptic markers (SNAP-25, a presynaptic membrane protein, and SYT-1, a synaptic vesicle protein). Figure 2 illustrates the different associations found between PTPRS isoforms (Q13332-2, −4, and −7) and SYT1 and SNAP25 in the frontal cortex. SYT1 protein levels negatively correlated with PTPRS type 2 (R^2^=0.12, P < 0.001), type 4 (R^2^=0.10, P <0.001), and type 7 (R^2^: 0.12, p <0.001) (Fig. 2, A and B). In contrast, levels of SNAP25 positively correlated with type 2 (R^2^=0.16, P < 0.001), type 4 (R^2^=0.11, P < 0.001) and type 7 (R^2^=0.05, P < 0.001) isoforms of PTPRS (Fig. 2, B). These associations suggest that PTPRS protein levels are linked to synaptic integrity and/or abundance in the CNS.

**Fig. 2:**
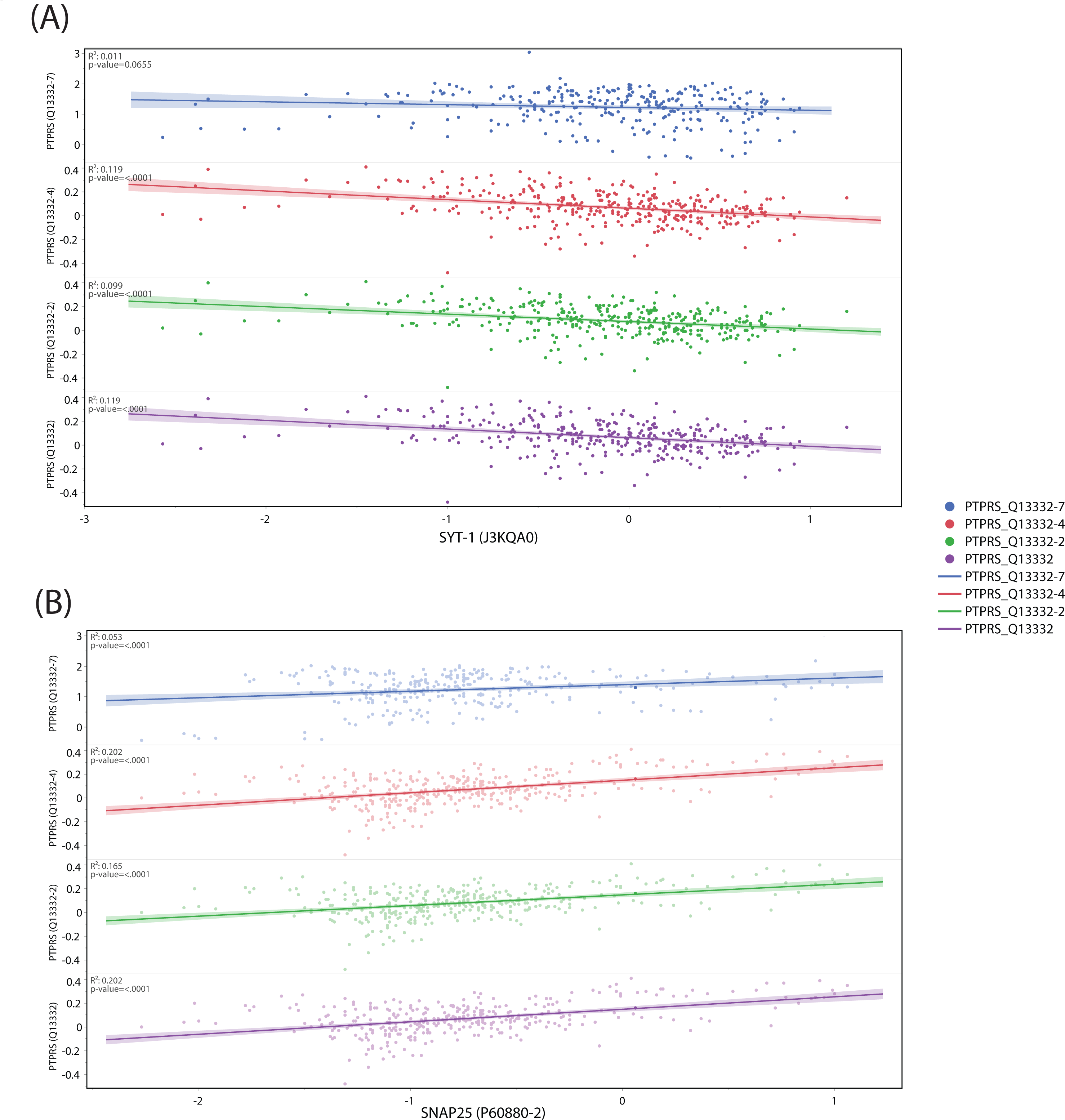
Correlations between PTPRS protein abundance and synaptic markers SYT1 and SNAP25. (A) Correlation between the abundance of four PTPRS protein isoforms (Q13332-7, Q13332-4, Q13332-2 and Q13332) and pre-synaptic marker SYT1 (J3KQA0). (B) Correlation between the abundance of four PTPRS protein isoforms (Q13332-7, Q13332-4, Q13332-2, and Q13332) and post-synaptic marker SNAP25 (P60880-2). Data extracted for the ROSMAP study. N(total)=360. Statistical analysis: (A and B) Simple linear regression with Pearson’s correlation (R^2^). Exact p-values are reported directly in the figures.

### Effect of *PTPRS* rs10415488 C variant on CSF P-tau levels in asymptomatic subjects with a parental history of AD, the PREVENT-AD cohort

We next sought to determine whether the presence of the rs10415488 C variant influenced AD biomarkers in pre-symptomatic but “at risk” individuals. To do so, we used biological samples from our PREVENT-AD cohort. Baseline CSF P-tau levels were significantly lower (p < 0.005) by the presence of the protective C allele in PREVENT-AD subjects (Fig. 3, A). A similar reduction was found when contrasting CSF total tau (T-tau)/Aβ42 ratios in *PTPRS* rs10415488 C carriers versus noncarriers (p < 0.05) (Fig. 3, B). A growing number of reports have also highlighted the link between neuronal macroautophagy and neurofibrillary tangles (29–31). Given the putative role of PTPRS in inhibiting autophagy, we sought to identify whether levels of the autophagic marker LC3 were changed between rs10415488 C carriers and non-carriers. We found that CC/CT carriers, who have a higher cortical abundance of PTPRS, had lower levels of LC3 compared to TT carriers (Fig. 3, C). Overall, these associations reveal that carriers of the C allele have reduced amounts of neurofibrillary tangles and presumably, healthier entorhinal cortices.

**Fig. 3:**
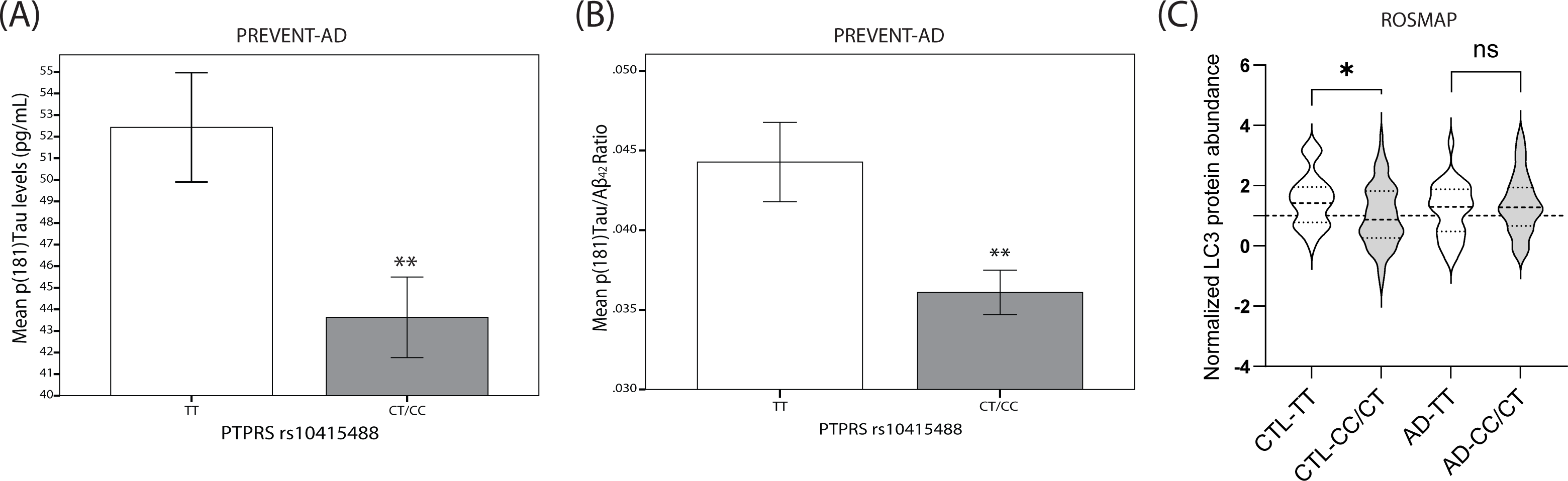
Lower CSF P-tau181 concentration in *PTPRS* rs10415488 C variant carriers in PREVENT-AD subjects. (A) Mean CSF P-tau181 concentration (pg/mL) in PREVENT-AD rs10415488 CT/CC carriers. N=36(TT) and 59(CT/CC). (B) Mean CSF P-tau181 concentration (pg/mL) normalized to Aβ42 levels in PREVENT-AD rs10415488 CT/CC carriers. N=36 (TT) and 59 (CT/CC). (C) LC3 protein abundance normalized to control (CTL) CC/CT carriers in the ROSMAP cohort. N=29(CTL-TT), 80(CTL-CT/CC), 30(AD-TT) and 58 (AD-CT/CC). Statistical analysis (A and B) Bilateral unpaired T-test (C) One-way ANOVA with Bonferroni’s multiple comparison and correction. ns= non-significant, *p≤0.05, **p≤0.01, ***p≤0.001, ****p≤0.0001.

### Effect of *PTPRS* rs10415488 C variant on entorhinal cortex volume levels and sex differences in CSF PTPRS protein levels in asymptomatic PREVENT-AD subjects

Using the same cohort, we next quantified the volume of the entorhinal cortex in carriers and non-carriers of the rs10415488 C variant. The volume of the entorhinal cortex is used as a surrogate measure for the severity of the disease since it is the brain region that is first affected in AD (32). Baseline entorhinal cortex volume (adjusted for intracranial cavity) was significantly increased (p < 0.05) in the presence of the protective C allele relative to non-carriers in asymptomatic but “at-risk” PREVENT-AD subjects (Fig. 4, A). We also found that the mean CSF PTPRS protein levels are markedly different between men and women (p<0.001), with women displaying a 40% reduction relative to men at baseline (Fig. 4, B).

**Fig. 4:**
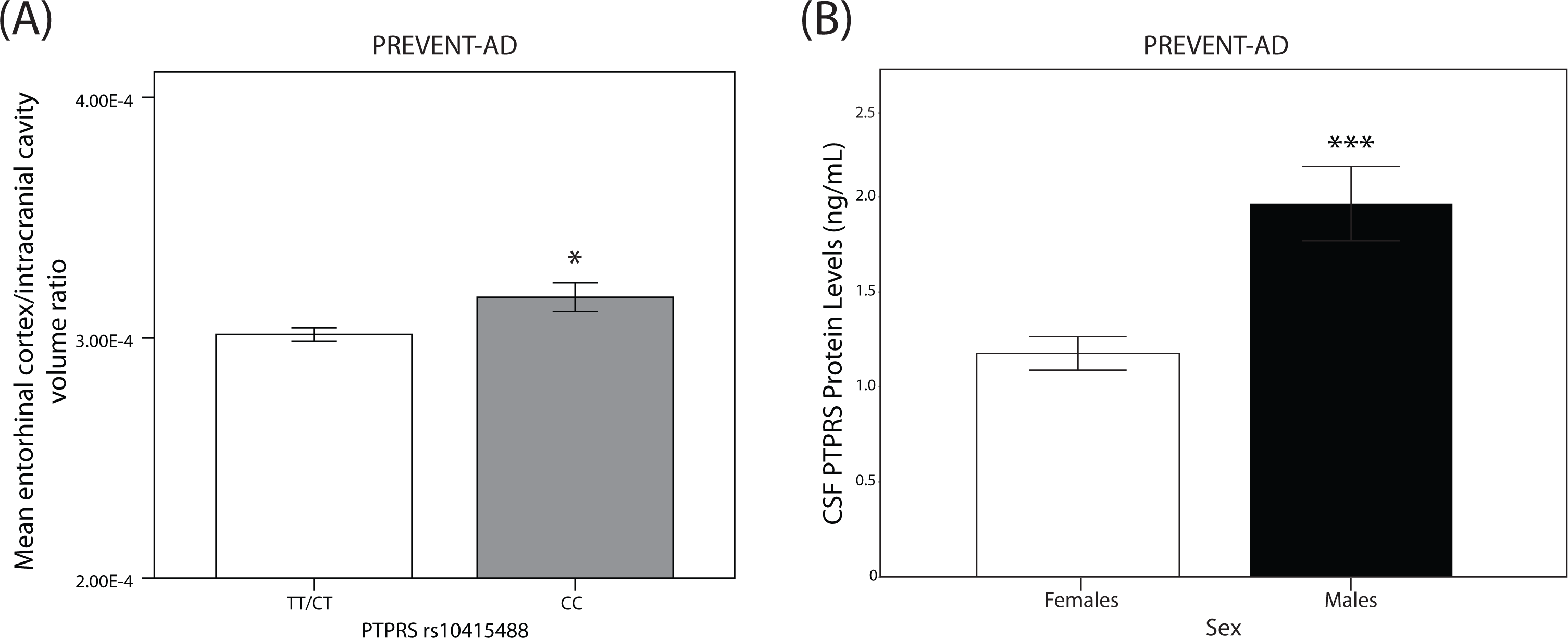
Increase entorhinal cortex volume in *PTPRS* rs10415488 homozygous CC carriers. (A) Mean entorhinal cortex volume normalized to intracranial cavity volume in PREVENT-AD rs10415488 TT/CT versus CC carriers. N=236(TT/TC), 43(CC). (B) Mean PTPRS protein abundance in the CSF of female and male PREVENT-AD subjects (ng/mL). N=71 and 31, respectively for females and males. Statistical analysis: (A and B) Bilateral unpaired T-tests. *p≤0.05, **p≤0.01, ***p≤0.001, ****p≤0.0001.

### Associations between CSF PTPRS protein levels and AD pathological biomarkers in *APOE4*-stratified PREVENT-AD subjects

Figure 5 illustrates the associations between CSF PTPRS protein levels and key AD pathological biomarkers [P-tau, T-tau, Aβ_42_, T-tau/Aβ42 ratio] following stratification by *APOE4*. Only T-tau (R^2^=0.08, P < 0.05) and P-tau (R^2^=0.06, P < 0.05) displayed significant associations in *APOE4*-negative subjects.

**Fig. 5:**
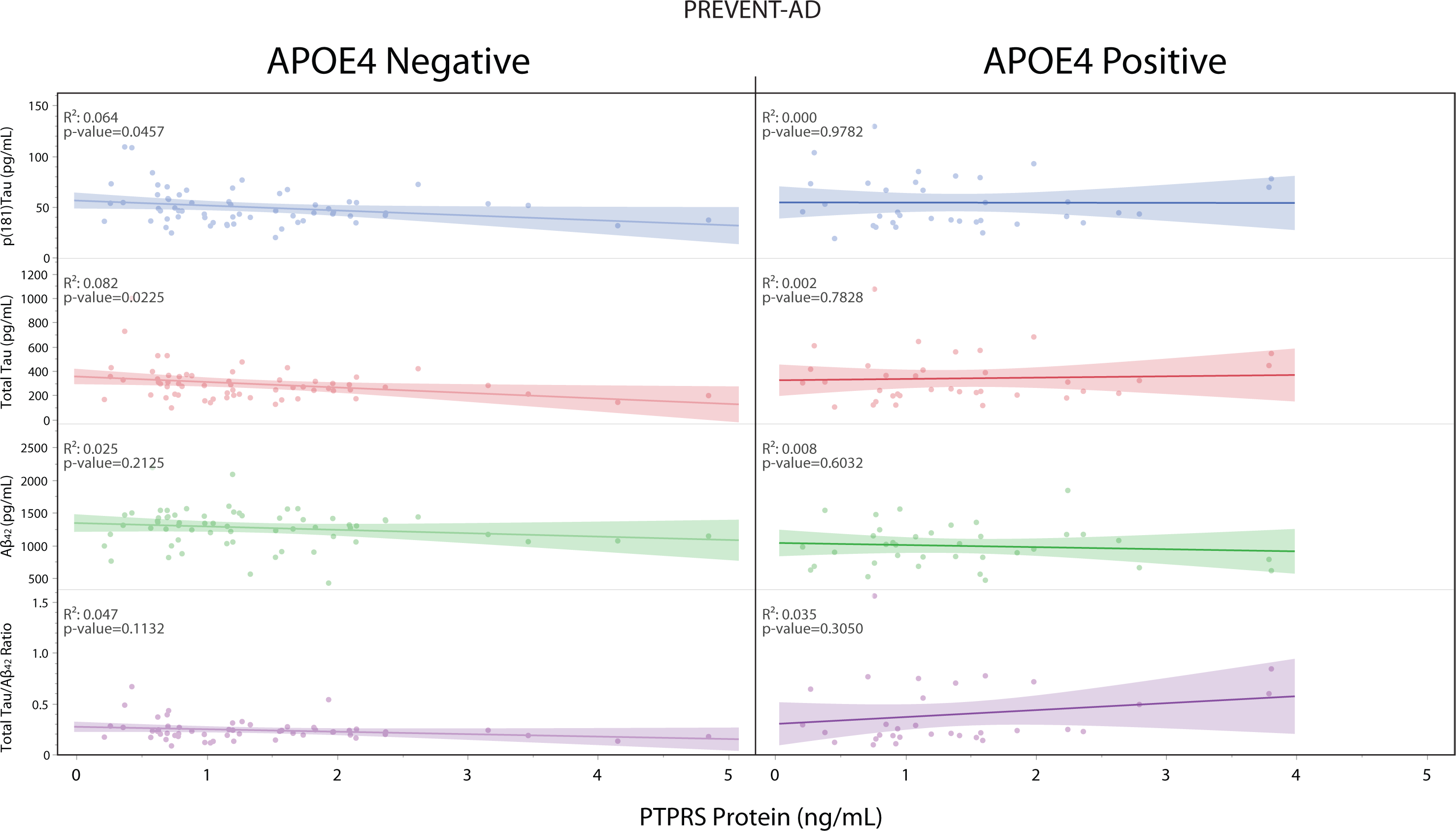
Correlations between AD biomarkers and the concentration of PTPRS in the CSF of PREVENT-AD *APOE4* carriers and non-carriers. PTPRS protein concentration (pg/mL) versus P-tau181, T-tau, Aβ42, and T-tau/Aβ42 levels in the CSF of PREVEN-AD subjects, stratified by *APOE4* status. N= 57 and 37 for *APOE4*(-) and (+) respectively. Statistical analysis: (A and B) Simple linear regression with Pearson’s correlation (R^2^). Exact p-values are reported directly in the figures.

### Protective *PTPRS* rs10415488 C variant is associated with reduced soluble GAP43 protein levels in the CSF and delayed AD onset in males

CSF levels of the axonal growth-associated protein GAP-43 were found to be reduced in carriers of the protective *PTPRS* rs10415488 C variant (Fig. 6, A). This effect was found to be restricted to the male population (p < 0.001), as no difference was observed in CSF GAP-43 levels in the presence or absence of the protective variant in females (Fig. 6, B). Finally, we investigated whether the presence of the protective C allele was linked to delayed AD onset using the ROSMAP study. We found that in male carriers, positivity for the C allele delayed the onset of AD by approximately ten months. These data showcase the link between PTPRS and synaptic integrity in males who are susceptible to AD.

**Fig. 6:**
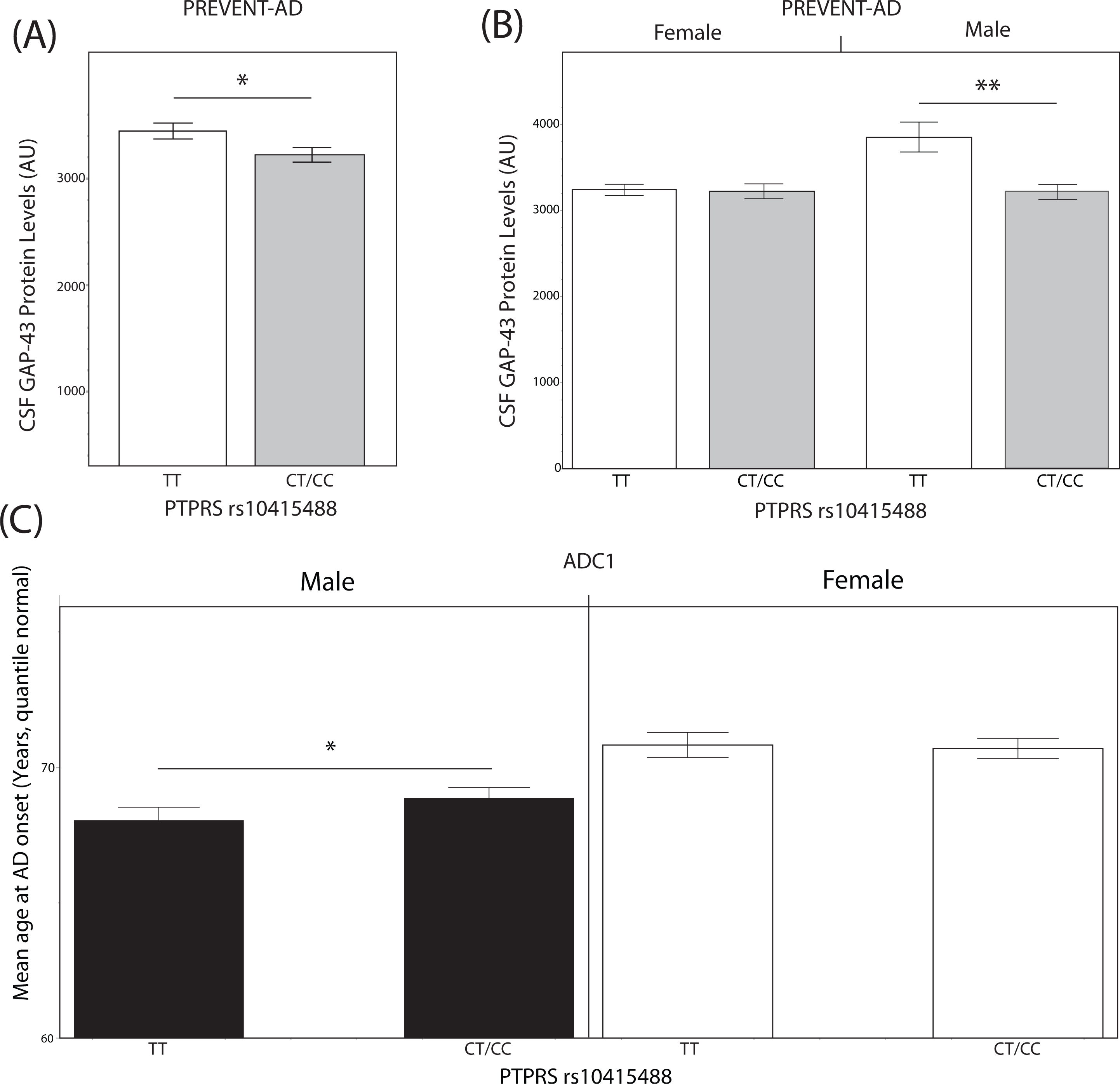
Association between PTPRS levels and synaptic integrity in PREVENT-AD subjects. **(A)** Mean CSF GAP-43 protein concentration in PTPRS rs10415488 C variant non-carriers and carriers. N=24 and 43 for TT and CT/CC carriers, respectively. **(B)** Sex stratification from (A). N=16, 31, 8, and 12 (from left to right). **(C)** Mean age at AD onset, stratified by sex in the ADCI cohort. N=321, 536, 344, and 586 (from left to right). Statistical analysis: (A-C) Bilateral unpaired T-tests. *p≤0.05, **p≤0.01, ***p≤0.001, ****p≤0.0001.

## Discussion

Studies on the roles of PTPRS and PTPRD in the central nervous system had initially focused on axon outgrowth, guidance, and reinnervation (33, 34). More recently, however, many cell biology studies have demonstrated novel roles of RPTPs as presynaptic proteins that trans-synaptically interact with multiple postsynaptic partners to mediate synaptic adhesion and synapse organization (2, 7, 35, 36).

Synaptic cell-adhesion molecules such as PTPRS form trans-synaptic complexes that are thought to initiate and maintain CNS plasticity (11, 37). They are also highly expressed in mature neurons, where they are believed to be essentially presynaptic (38). Cell-adhesion interactions of PTPRS with numerous postsynaptic partners, including NGL-3, TrkC, SALMs, SliTrks, and IL1RAPs, suggest a major role for RPTPs in synapse formation and/or remodeling (37, 39).

Our study reports significant associations between PTPRS and synaptic markers in both cortical tissues and CSF throughout the Alzheimer spectrum. Nevertheless, this was found without association with classical AD pathological markers such as neurofibrillary tangles and senile plaques (Fig 1, C, and D). When *APOE4* stratification was considered, slight changes were noticed for the CSF tau biomarkers P-tau and T-tau in *APOE4*-negative PREVENT-AD subjects (Fig. 5). On the other hand, tissue PTPRS mRNA levels are decreased in cortical areas of AD patients compared to controls in *APOE4*-positive subjects, when the end-stage brains are saturated with both amyloid and tau deposits. The high levels of both CSF P-tau and T-tau protein in asymptomatic subjects expressing low levels of PTPRS are certainly consistent with early signs of tau pathology (P-tau) and neuronal degeneration (T-tau) in the absence of visible (or detectable) deposition on PET scans (data not shown).

To examine further the relationship between PTPRS and AD, we examined the genetic contribution of the *PTPRS* gene locus. A first step we identified by eQTL, a polymorphism, rs10415488, which affects PTPRS expression in an allele-dependent manner and provides some level of protection in two different case/control populations from North America and Europe (Fig. 1, A and Table I). The presence of the C allele is associated with higher expression levels in the frontal, occipital, and temporal cortices and in the putamen of normal subjects devoid of any neuropathologies (Fig. 1, A). The risk reduction level observed in the QFP population isolate of eastern Canada (OR = 0.86, p < 0.05) is more pronounced than the one observed with the large heterogeneous IGAP GWAS cohort (OR =0.97, 0 < 0.05). Two of the main characteristics of the QFP that set it apart are that all subjects are descendants of a group of 3000 migrants that came to North America about 400 years ago. It thus forms a very homogeneous population with little genetic background variability. Furthermore, cases and controls were matched for age, sex, and most importantly, place of birth, an adjustment rarely performed in GWAS studies. In other words, the C allele is associated with high expression levels of PTPRS and a lower risk of AD.

When we transposed these observations to our pre-symptomatic “at-risk” asymptomatic subjects, we found that the protective allele C is associated with significantly lower levels of CSF P-tau in carriers (p <0.004) and a corresponding reduction in the P-tau/Aβ42 ratio in the same subjects (Fig. 3). The latter ratio commonly serves as an indicator of AD neuropathology, which is associated with the short-term emergence of cognitive deficits in asymptomatic subjects. In addition, analysis of entorhinal cortex volume by structural MRI revealed significantly higher cortical volume in protective rs10415488 C allele carriers when compared to TT subjects. The entorhinal cortex volume decline is typical of normal aging and was shown to be accelerated by subjects exhibiting cognitive deficits and AD. It is also considered by many as the starting point of tangles deposition and spreading in sporadic AD.

One molecular function of PTPRS we chose to explore was macroautophagy, given the implication of this biological process in the response against neurodegeneration. In neurons, it was shown that LC3 immunoreactivity occurred in most dystrophic neurites in AD and co-localized with abnormal P-tau in many neurofibrillary tangles (40). Co-localization of autophagy markers with P-tau has been observed previously in PS1/APP transgenic mice (41). In cell culture models of AD, autophagy markers are induced prior to accumulation of phosphorylated tau (30). In the AD brain, these markers of neuronal autophagy dominate early in the disease and decline with increasing disease duration, indicating their expression in neurons targeted for neurodegeneration over time (Fig. 3, C) (40).

The data generated from this study is dichotomous to previously published mouse models and human brain datasets. Indeed, previous knock-out mouse studies suggested that deficiency of PTPRS had a protective effect on neuronal plasticity (7, 14, 42). These seemingly opposite results are plausibly the consequence of using acute spinal cord injury as a model for exploring the function of PTPRS. Furthermore, we are limited by modest changes in the total abundance of PTPRS. Perhaps a complete knockout of the gene gives rise to compensation mechanisms. Given that PTPRS orchestrates numerous biological functions, it is thus likely that the enzyme operates differently in the context of acute injury than it does during chronic affliction, such as in AD.

We also highlighted the presence of a sexual dimorphism in the protein abundance of PTPRS. Given the putative protective role of PTPRS in human AD pathology, the lower abundance of the phosphatase in women might impact the risk of developing or the severity of AD. This result goes in line with the fact that AD disproportionately affects women. The sex differences observed in CSF PTPRS protein levels with significantly higher levels observed in men is intriguing (Fig. 4, B), especially when paired with reduced GAP-43 abundance in CSF (Fig. 6, C). In contrast, women, which seem to constitutively express less PTPRS than men in the CNS, are not protected by the rs10415488 C variant. These differences are perhaps the result of sex-based regulatory targeting patterns for PTPRS expression, but that remains conjecture.

## Conclusions

Dose of rs10415488 variant C increases the expression of PTPRS in the brain, which in turn acts as a protective factor for early tau pathology in AD. PTPRS protein levels were found to be higher in males than females, with variant C significantly delaying the age of onset of AD in males. These observations support retrospective evidence linking higher PTPRS levels to decreased risk of developing AD. Overall, this work highlights a protective role for PTPRS in the asymptomatic phase of AD and sediments the crucial role of R2B phosphatases in neurodegeneration and synaptic plasticity.

## Declarations

### Acknowledgements

We wish to thank Jennifer Tremblay-Mercier, Doris Dea, and Louise Théroux for their individual contribution at different stages of the project. We also thank Dr. Naguib Mechawar at the Douglas Institute/ Bell Canada Brain Bank for providing human brain tissues from the Québec Founding Population.

### Authors contributions

AP, JP, DA, MT, CP, HZ, and KB conceptualized the research. AL, IA, and members of the PREVENT-AD research group performed CSF biomarkers measurements, data quality control, and data compilation. CP performed the QTL analyses of DNA samples and risk assessments. AP, JP, CP, and MT contributed to the data analysis. AP, JP, and MT wrote the original manuscript draft. All authors reviewed, edited, and approved the final manuscript.

### Ethical approval and consent to participate

All procedures were approved by the McGill University Faculty of Medicine Institutional Review Board and complied with the ethical principles of the Declaration of Helsinki. In the QFP cohort, 382 individuals were genotyped and selected for evaluation. Each participant and study partner provided written informed consent.

### Consent for publication

All authors have approved the content of this manuscript and provided consent for publication.

### Availability of data and materials

All the data necessary for raising the conclusions are found within the manuscript and the supplemental files.

### Funding

AP is a Canderel Studentship and FRSQ doctoral scholarship recipient. ML Tremblay is supported by the J. and J.L. Levesque Chair in Cancer Research, a Canadian Institute of Health Research Foundation grant to (CIHR FDN-159923), the Richard and Edith Strauss Canada Foundation, and the Aclon Foundation. J Poirier is supported by the Fonds de la Recherche en Santé du Québec (FRSQ), the Canadian Institute for Health Research (CIHR # PJT 153287), and the J.L. Levesque Foundation. Dr. Zetterberg is a Wallenberg Scholar supported by grants from the Swedish Research Council (#2022-01018 and #2019-02397), the European Union’s Horizon Europe research and innovation programme under grant agreement No 101053962, Swedish State Support for Clinical Research (#ALFGBG-71320), the Alzheimer Drug Discovery Foundation (ADDF), USA (#201809-2016862), the AD Strategic Fund and the Alzheimer’s Association (#ADSF-21-831376-C, #ADSF-21-831381-C, and #ADSF-21-831377-C), the Bluefield Project, the Olav Thon Foundation, the Erling-Persson Family Foundation, Stiftelsen för Gamla Tjänarinnor, Hjärnfonden, Sweden (#FO2022-0270), the European Union’s Horizon 2020 research and innovation programme under the Marie Skłodowska-Curie grant agreement No 860197 (MIRIADE), the European Union Joint Programme – Neurodegenerative Disease Research (JPND2021-00694), the National Institute for Health and Care Research University College London Hospitals Biomedical Research Centre, and the UK Dementia Research Institute at UCL (UKDRI-1003). Dr. Blennow is supported by the Swedish Research Council (#2017-00915), the Alzheimer Drug Discovery Foundation (ADDF), USA (#RDAPB-201809-2016615), the Swedish Alzheimer Foundation (#AF-742881, #AF-930351, #AF-939721 and #AF-968270), Hjärnfonden, Sweden (#FO2017-0243 and #ALZ2022-0006), the Swedish state under the agreement between the Swedish government and the County Councils, the ALF-agreement (#ALFGBG-715986 and #ALFGBG-965240), the European Union Joint Program for Neurodegenerative Disorders (JPND2019-466-236), the National Institute of Health (NIH), USA, (grant #1R01AG068398-01), and the Alzheimer’s Association 2021 Zenith Award (ZEN-21-848495). Funds from the Victor Phillip Dahdaleh Institute of Genomic Medicine at McGill University supported the genotyping.

Data used in preparation of this article were obtained from the program of PRe-symptomatic EValuation of Novel or Experimental Treatments for Alzheimer’s Disease (PREVENT-AD) at the Centre for Studies on Prevention of Alzheimer’s Disease (StoP-AD), Douglas Mental Health University Institute Research Center (https://preventad.loris.ca/main.php). A complete listing of the PREVENT-AD Research Group can be found at: https://preventad.loris.ca/acknowledgements/acknowledgements.php?date=2023-09-21.

**<u>Consortia</u>**

**PREVENT-AD research group**

Judes Poirier^17,18,19^, John C. S. Breitner^17,18^, Alexandre Poirier^17^, Justin Miron^17^, Cynthia Picard^17^, Anne Labonté^17^, Sylvia Villeneuve^17,18,19^, R. Nathan Spreng^19^, Pedro Rosa-Neto^18^ & Jennifer Tremblay-Mercier^17^.

^17^Douglas Mental Health Institute Research Centre, McGill University, Montreal, Quebec, Canada

^18^Department of Psychiatry, McGill University, Montreal, Quebec, Canada

^19^Department of Neurology and Neurosurgery, McGill University, Montreal, Quebec, Canada

## Abbreviations

Aβ: amyloid beta
Aβ_42_: amyloid beta 42 amino acid peptide
AD: Alzheimer’s disease;
ADC1: Alzheimer Disease Center Dataset 1
*APOE4*: apolipoprotein E (ε4 allele)
BRAINEAC: Brain eQTL Almanac
CERAD: Consortium to Establish a Registry for Alzheimer’s Disease
CNS: central nervous system
CSF: cerebrospinal fluid
CTL: controls
DNA: deoxyribonucleic acid
ELISA: enzyme-linked immunosorbent assay
eQTL: expression quantitative trait loci
FCTX: Frontal cortex
HIPP: Hippocampus
LC/MS/MS: liquid chromatography and mass spectroscopy
MCI: mild cognitive impairment
mRNA: messenger ribonucleic acid
PREVENT-AD: pre-symptomatic evaluation of experimental or novel treatments for Alzheimer’s disease
P-tau: Tau protein phosphorylated at amino acid residue Threonine 181
RNA: ribonucleic acid
ROSMAP: Religious Orders Study, and the Memory and Aging Project
SNP: single nucleotide polymorphism
t-tau: total-tau protein

