## Supplemental Figures for "PTPRS is a novel marker for early tau pathology and synaptic integrity in Alzheimer’s disease"

**Supplementary materials:**
Supplemental Figures:

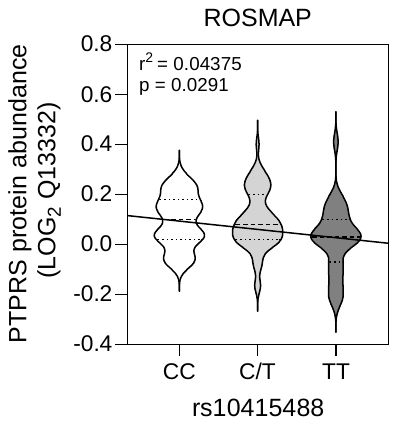

**Sup. Fig. 1: rs10415488 variant T allele status is negatively correlated with the protein abundance of PTPRS in the frontal cortex in the ROSMAP cohort**

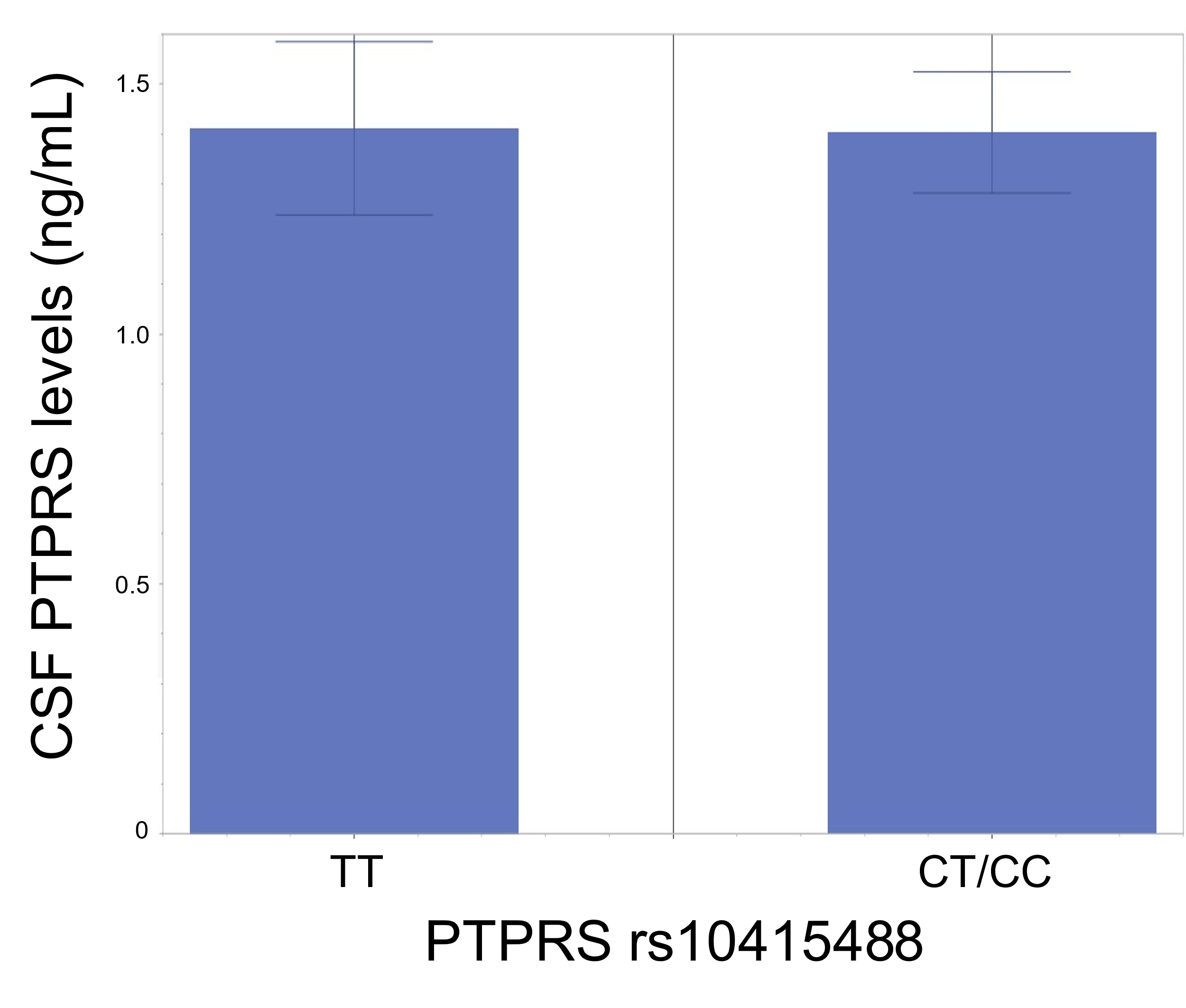

**Sup. Fig. 2: rs10415488 variant status does not affect PTPRS protein concentrations in the CSF**

The concentration of soluble PTPRS between non-carriers (TT) or carriers (CT/CC) (ng/mL) in the CSF of PREVENT-AD subjects.

| **Cohort** | **Stratification**  **APOE4** | **Major Allele**  **(Homozygous)** | **N AD/CTL** | **OR** | **p-val.** |
| --- | --- | --- | --- | --- | --- |
| Quebec Founding Population  (Homogenous European) | -- | TT | 964/984 | 1.30 | 0.008 |
| QFP* | E4-Positive  E4-Negative | TT  TT | 476/176  486/808 | 3.12  1.22 | 0.05  0.04 |
| AD Centres Consortium  (heterogenous) | -- | TT | 1981/509 | 1.18 | 0.09 |
| ADCC-ADC1** | E4-Positive  E4-Negative | TT  TT | 1252/130  729/379 | 0.93  1.37 | n.s.  0.02 |

**Sup. Table I: rs10415488 variant TT is a risk allele for Alzheimer’s disease in multiple cohorts**

*Quebec founding population datasets is available at https://www.PREVENTAD.loris.ca and **ADCC-ADC1 genotypes information is available at <https://www.niagads.org/datasets/ng00022> for registered investigators.
